# Comparing optimal transport and machine learning approaches for databases merging in scenarios involving missing data in covariates. Application to Medical Research

**DOI:** 10.64898/2026.01.23.701369

**Authors:** Flore N’kam Suguem, Sébastien Déjean, Philippe Saint Pierre, Nicolas Savy

**Affiliations:** Institut de Mathématiques de Toulouse, UMR521, Université de Toulouse, CNRS, France; Université de Toulouse UT, F-31062 Toulouse Cedex 9, France; Université Toulouse Jean-Jaurès UT2J, F-31058 Toulouse, France; Department of Physics and Astronomy, University of Bologna, 40127 Bologna, Italy

**Keywords:** optimal transport, data integration, missing data, statistical matching, data recoding

## Abstract

**Motivation:** One of the challenges encountered when merging heterogeneous observational clinical datasets is the recoding of categorical target variables that may have been measured differently across data sources. Standard machine learning-based approaches, such as Multiple Imputation by Chained Equations and the k-Nearest Neighbours method are compared with an Optimal Transport based algorithm (OTrecod) when databases are altered by missing values in covariates or by imbalanced groups. The empirical performance in these realistic data integration settings remains underexplored.

**Results:** A comprehensive simulation study was conducted, varying sample size, group imbalance, signal-to-noise ratio, and mechanisms of missing data. The results demonstrate that OTrecod consistently achieves higher recoding accuracy compared with Multiple Imputation by Chained Equations and k-Nearest Neighbours, particularly in large, imbalanced and weak-signal scenarios. These findings are further illustrated using subsets of the National Child Development Study, where OTrecod and Multiple Imputation by Chained Equations minimised the distributional divergence between recoded social-class scales, while k-Nearest Neighbours produced less stable results.

**Availability and Implementation:** The source code supporting this study is publicly available at https://github.com/FloAI/CompareOT.

## 1 Introduction

Today, open science is becoming increasingly prevalent in the production and dissemination of scientific knowledge. This approach is based on the transparent sharing of data, methods, and results throughout the research process. Researchers are thus encouraged to make their data openly available to the scientific community. Finding data is thus becoming easier. But finding data is not enough, and in this context, the Findability, Accessibility, Interoperability, and Reuse (FAIR) principles provide guidelines to go beyond simply finding data and extends to effective reuse and integration. To make these guidelines efficient, robust methodological developments are essential. In particular, data interoperability often involves integrating heterogeneous databases, originally collected for different purposes. Our contribution is situated within this broader challenge. More specifically, improving database merging, a key step in enabling interoperable and reusable scientific data is the motivation for our work.

Different techniques are already used to combine heterogeneous databases from diverse sources, leveraging both statistical and symbolic approaches. These include probabilistic models and probabilistic databases through principled probabilistic reasoning (Suciu, 2020), time-dependent or sequential data through the use of Hidden Markov Models and temporal models employed to align, smooth, and infer latent states across data streams, supporting robust temporal integration (Castanedo, 2013). At a higher semantic level, other techniques are used such as logical reasoning techniques such as ontology-based integration, rule engines, and schema mapping help reconcile semantic heterogeneity and ensure interoperability between differing data representations (Borowicc and Souza, 2024; Putrama and Martinek, 2024). More recently, knowledge graph frameworks have emerged for improved scalability and interpretability. Further extensions of these paradigms through hybrid AI approaches combine large language models and graph learning with probabilistic and logical reasoning to automate schema discovery, data fusion, and knowledge validation across heterogeneous systems (Mohamed et al., 2025).

One main challenge that arises in database integration is that of variable recoding. This problem emerges especially when a categorical variable of interest is represented using different coding schemes across two different databases. Such inconsistencies, often stem from differences in data collection or interpretation methods, lead to significant difficulties in the integration process. We look at Database A containing covariates and one version of the target variable *Y*_*A*_. Database B contains an alternative encoding *Y*_*B*_ of the target variable *Y*. The goal is to create a synthetic database for *Y*_*B*_ in database A and *Y*_*A*_ in database B (Gares et al., 2019). There is an important literature dealing with this question. A natural approach is to consider imputation methods (Rubin, 1987) such as the Multiple Imputation by Chained Equations (MICE) (van Buuren and Groothuis-Oudshoorn, 2011) to impute the missing values in *Y*_*A*_ and *Y*_*B*_. Another approach consist in predicting the missing values in *Y*_*A*_ and *Y*_*B*_ using statistical or supervised learning models (Jang et al., 2020; Alwateer et al., 2024). Finally, one can also consider approaches based on optimal transport aiming to exploit the transportation plan between the probability distributions *Y*_*A*_ and *Y*_*B*_ (Gares et al., 2019; *Garès and Omer, 2020)*.

The aim of the paper is to evaluate the performance of the three previous approaches in this variable recoding context arising in data merging. In particular, we focus on exploring the effects of different types of data biases on the performance of the methods. Potential sources of bias include the amount and mechanism of missing data in covariates, correlations between covariates and outcomes or distributional shifts of the variable of interest. One of the most critical factors when addressing such recoding problem is that of missing data in the covariates. This poses a fundamental challenge in statistical analysis and machine learning, as it can introduce bias, reduce statistical power, and compromise the validity of conclusions if not properly addressed (Little, 2024; Rubin, 1987). Such issues occur across multiple domains, including healthcare, finance, and social sciences, where incomplete databases often result from measurement errors, data collection constraints, or participant non-response (Schafer, 1997; van Buuren, 2018). To address this issue, imputation techniques such MICE van Buuren and Groothuis-Oudshoorn (2011) are widely used. In this paper, a simulation study is performed to evaluate the effect of data bias due to correlation, distributional shift and missing data (amount of missing data, missing data management, mechanism of missing data).

Background on the three approaches used for recoding problem and on missing data mechanisms are given in Section 2. Section 3 introduces the simulation study and the various steps used to simulate the database. Different scenarios are proposed to evaluate the performance of each approaches. Results of the simulation study are given and discussed in Section 4. Each approaches are also evaluated on a real-world database, the National Child development Study (NCDS), to assess their practical applicability 5. Section 6 concludes the paper and discuss futures works.

## 2 Background

### 2.1 Methods for recoding variables to merge databases

#### 2.1.1 Algorithm OT based on optimal transportation

The idea behind the OT algorithm is to exploit an optimal transportation plan between the probability distributions of Y_A_ and Y_B_. Optimal Transport (OT) theory, originally introduced by Gaspard Monge in 1781 (Monge, 1781), aims at finding a measurable mapping that reallocates one distribution of mass onto another while minimising a prescribed transportation cost. In the continuous setting, under suitable regularity assumptions on the cost function (e.g., strict convexity), the optimal transport plan is characterised by a unique transport map that can often be expressed as the gradient of a convex potential (Brenier’s theorem). In the categorical or discrete setting, the transport plan is instead formulated as a coupling matrix solving a linear programming problem, commonly referred to as the Hitchcock’s transportation problem; in that case, the solution is generally not unique but can be efficiently computed using dedicated optimisation algorithms.

Given two databases *A* and *B*, where *Y*_*A*_ and *Y*_*B*_ are respectively observed, one can estimate an empirical optimal transport plan by defining a cost function between individuals based on the covariates that are jointly observed in both datasets. This principle underlies the recoding and data fusion approaches proposed in (Gares et al., 2019; Garès and Omer, 2020), where regularised optimal transport is used to construct a probabilistic linkage between units from the two samples through a cost reflecting their similarity in the common covariate space. The empirical optimal transport plan give us the estimated number of individuals recoded in each modality of the variable to be recoded. The second step of allocation of a given individual to a modality is done by means of k-nearest neighbours thanks to the cost function introduced.

More broadly, recent methodological and computational advances have established OT as a powerful framework for modern data analysis and integration. It provides a principled way to align and compare probability distributions arising from heterogeneous sources, with applications ranging from domain adaptation to data fusion and covariate shift correction (Courty et al., 2017; Peyré and Cuturi, 2019). These developments are accompanied by practical implementations, such as the OTrecod R package, which makes OT-based data fusion approaches readily accessible for applied statistical analysis (Guernec et al., 2023).

#### 2.1.2 Multivariate Imputation Chained Equations (MICE)

The Multiple Imputation by Chained Equations (MICE) algorithm represents a major advancement within the Multiple Imputation framework (van Buuren and Groothuis-Oudshoorn, 2011). MICE adopts a fully conditional specification strategy. It iteratively imputes each incomplete variable using a regression model conditioned on all other variables. This chained procedure allows for flexible model specification. MICE is a powerful tool for database integration as it uses convergence to complete databases.

Each variable with missing data is modelled conditionally upon the other variables in the dataset as part of the MICE process, which involves running a number of regression models. This makes it possible to represent every variable in accordance with its distribution. Logistic regression is used to model binary variables, while linear regression is used to model continuous variables (van Buuren and Groothuis-Oudshoorn, 2011).

There are four primary steps in the MICE process. Initially, basic placeholder values, like the mean or mode, are used to impute each variable with missing data. After that, all other values are retained, but the imputed values for one variable are returned to missing. This variable is then treated as the dependent variable and the other variables as independent predictors when it is regressed on the other variables in the dataset. The missing values for that variable are substituted with the predictions that are produced, frequently with additional random variation. The imputations can be improved one step at a time until they stabilise by repeating these procedures for every variable with missing data and cycling through several iterations. After that, a number of finished datasets are produced in order to capture the imputation process’s uncertainty.

MICE is widely used in applied research in the social sciences, psychology, and epidemiology because of its practical performance and methodological flexibility. Multivariate relationships have been well established through simulation and empirical research, and they preserve correlations between variables without the need for explicit distributional assumptions.

#### 2.1.3 k-Nearest Neighbours (kNN)

The K-Nearest Neighbours (kNN) algorithm is a non-parametric, instance-based learning method commonly used for both classification and regression tasks (Cover and Hart, 1967). The core principle of kNN is to predict the value or class of a given instance, the algorithm identifies the *k* most similar instances, or neighbours, in the feature space and bases its prediction on their values. In classification problems, kNN typically employs a majority voting scheme, while in regression, it uses the mean or median of the neighbours’ values. The similarity between instances is generally quantified through distance metrics such as the Euclidean or Manhattan distance. kNN is also used to solve problems in missing data imputation. Here, kNN relies on the assumption that similar samples possess similar feature values. In this case, the algorithm first identifies one or more missing attributes and their *k* nearest neighbours based on the observed non-missing features. The distances between its variables can be computed, typically through a normalised Euclidean distance to ensure all features contribute equally. For continuous variables, the imputed value is often the mean or median of the neighbours’ values, while in the case of categorical variables, the most frequent category is used.

### 2.2 Missing data

Various techniques can be employed to deal with missing data, broadly categorised into deletion methods and imputation techniques. Deletion methods, such as list-wise and pairwise deletion, remove incomplete cases from the dataset. Although simple to implement, these approaches can lead to substantial information loss and biased estimates (Rubin, 1976; Allison, 2002). Basic imputation techniques, such as mean or median imputation, are widely used but can distort variability and underestimate uncertainty (Donders et al., 2006). More advanced techniques, including multiple imputation (MI) (Rubin, 1987), expectation maximisation (EM) (Dempster et al., 1977), and machine learning-based imputation (Jang et al., 2020; Alwateer et al., 2024), offer more robust solutions by leveraging patterns in the data.

According to Rubin (1976) (Little, 2024), there are three missingness mechanisms: Missing Completely at Random (MCAR), where the probability of missingness is independent of both observed and unobserved data; Missing at Random (MAR), where missingness depends only on the observed data; and Missing Not at Random (MNAR), where missingness depends on the unobserved data itself.

## 3 Simulation Design

Performance of the different recoding methods under various scenarios are assessed by means of simulations. Based on a set of covariates, we generate a continuous outcome that we convert into a categorical one according to two discretisations with a different number of categories, to mimic two databases with different encoding of one variable *Y*. Then values in covariates are amputated to address the missing values issue. The different steps of the simulation design are illustrated in Figure 1 and formalised below.

**Figure 1.**
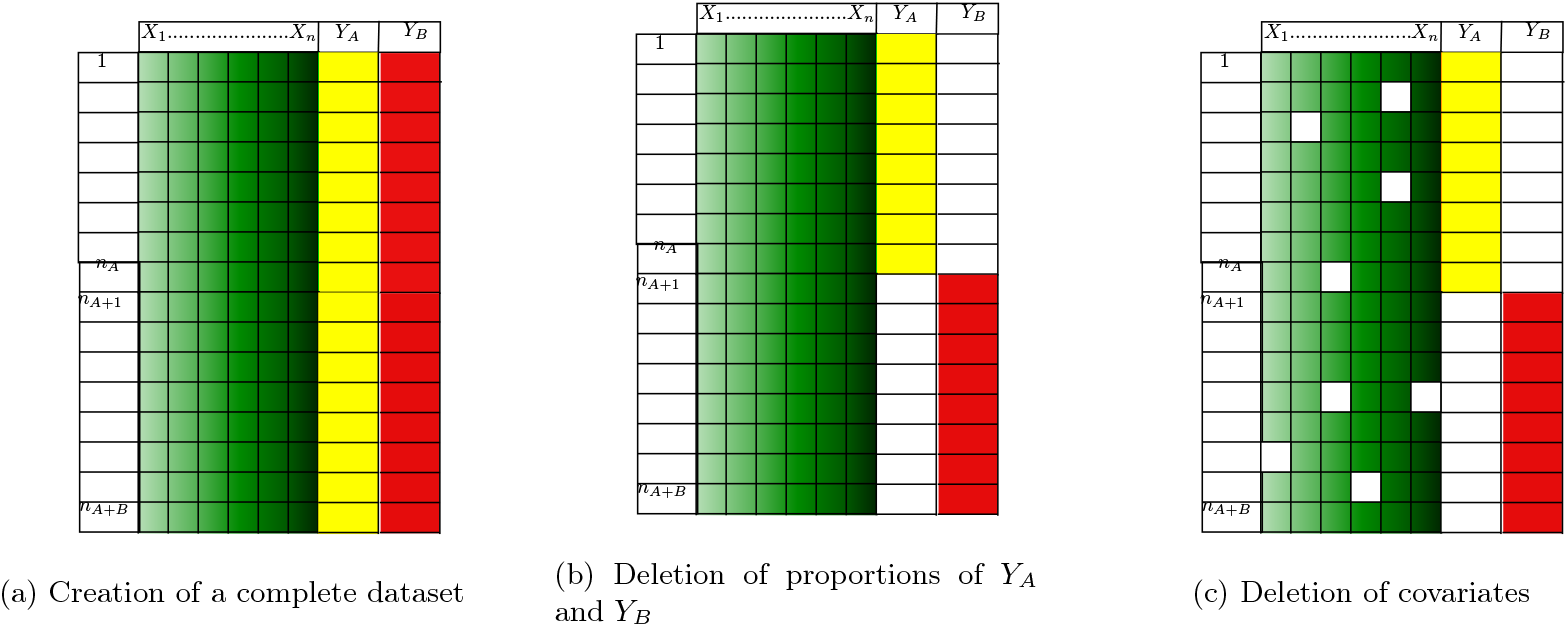
Workflow of dataset generation

### 3.1 Complete database generation

To generate the complete database, first we consider (*X*_1_, …, *X*_*p*_) a set of *p* independent covariates with fixed distributions. Then we construct *Z* a continuous outcome as a linear combination of the covariates perturbed by an independent Gaussian random noise:

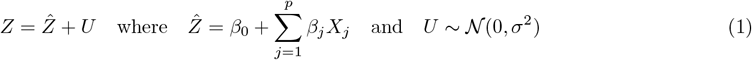

The strength of association between the covariates and the *Z* may be assessed by means of the coefficient of determination *R*^2^ which quantifies the proportion of variance in the outcome *Z* that is explained by the covariates and is defined as:

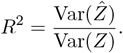

The distributions of the covariates being given, 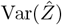 can easily be derived. Noticing that

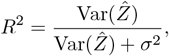

and thus

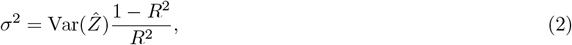

it is thus possible to fixed the strength of association between the covariates and the outcome by varying the values of *σ*^2^.

To investigate the impact of the assumption of equality of the distribution *Z*_*A*_ and *Z*_*B*_, two scenarios are considered:

- **Scenarios S (for “same”):** assumes *Z*_*A*_ and *Z*_*B*_ have the same distribution. To do so, the values of the coefficients in the definitions 1 are the same for *Z*_*A*_ and *Z*_*B*_.
- **Scenarios D (for “different”):** consider *Z*_*A*_ and *Z*_*B*_ have not the same distribution. To do so, the values of the coefficients in the definitions 1 are not the same for *Z*_*A*_ and *Z*_*B*_.

Finally, given *n*_*A*_ (resp. *n*_*B*_) the sample size of database *A* (resp. *B*), given a value of *R*^2^, and given a scenario S or D, the complete database generation follows the following steps:

1. Generate *n* = *n*_*A*_ + *n*_*B*_ values (*x*_1,*i*_, …, *x*_*p,i*_), *i* = 1, …, *n* through their distributions,
2. Generate *n* = *n*_*A*_ + *n*_*B*_ values *u*_*i*_, *i* = 1, …, *n* from a centred Gaussian distribution of variance *σ*^2^ given by relation (2).
3. For each *i* = 1, …, *n*, compute *z*_*A,i*_ and *z*_*B,i*_ by means of relation (1).
4. (*y*_*A,i*_, *i* = 1, …, *n*) is (*z*_*A,i*_, *i* = 1, …, *n*) categorised by quartiles and (*y*_*B,i*_, *i* = 1, …, *n*) is (*z*_*B,i*_, *i* = 1, …, *n*) categorised by tertiles.

Data generated at this step is illustrated by Sub-Figure 1a. The data used to test the performances of the algorithms (Sub-Figure 1b) consists in Database A : (*x*_1,*i*_, …, *x*_*p,i*_, *y*_*A,i*_, *i* = 1, …, *n*_*A*_), and Database B : (*x*_1,*i*_, …, *x*_*p,i*_, *y*_*B,i*_, *i* = *n*_*A*_ + 1, …, *n*_*A*_ + *n*_*B*_).

#### Remark 1

*The remaining values* (*y*_*A,i*_, *i* = *n*_*A*_ + 1, …, *n*_*A*_ + *n*_*B*_) *and* (*y*_*B,i*_, *i* = 1, …, *n*_*A*_) *are used to assess the performance of the algorithm (see Section 3.5)*.

In order to simulate realistic patterns resembling those observed in the National Child Development Study (NCDS) (see Subsection 5 for details), five covariates are considered with the distributions listed in Table 1. For scenario S, the coefficients are fixed to:

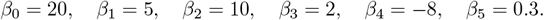

and for scenario D, they are fixed to

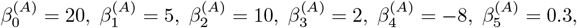

for *Z*_*A*_ and to

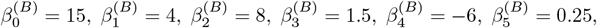

for *Z*_*B*_.

**Table 1.**
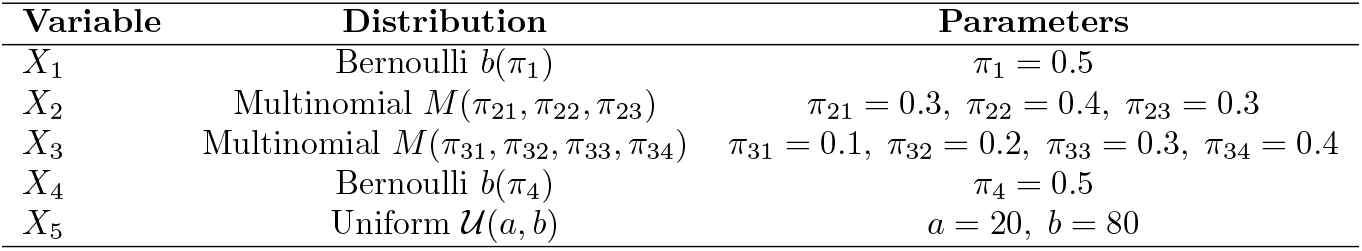
Distributions of the covariates for the simulation study.

| Variable | Distribution | Parameters |
| --- | --- | --- |
| $X_1$ | Bernoulli $b(\pi_1)$ | $\pi_1 = 0.5$ |
| $X_2$ | Multinomial $M(\pi_{21}, \pi_{22}, \pi_{23})$ | $\pi_{21} = 0.3, \pi_{22} = 0.4, \pi_{23} = 0.3$ |
| $X_3$ | Multinomial $M(\pi_{31}, \pi_{32}, \pi_{33}, \pi_{34})$ | $\pi_{31} = 0.1, \pi_{32} = 0.2, \pi_{33} = 0.3, \pi_{34} = 0.4$ |
| $X_4$ | Bernoulli $b(\pi_4)$ | $\pi_4 = 0.5$ |
| $X_5$ | Uniform $\mathcal{U}(a, b)$ | $a = 20, b = 80$ |

### 3.2 Introduction of missing data in the covariates

In order to investigate the behaviour of the algorithm when data missed in the covariates, a given proportion *P* of values ((*x*_1,*i*_, …, *x*_*p,i*_), *i* = 1, …, *n*) are randomly deleted according to the missing data mechanism chosen. The resulting datasets *A* and *B* is illustrated by Sub-Figure 1c. To generate missing data in the covariates (known as amputation procedures), R-miss-tastic package developed by Mayer et al. (2022) is used. R-miss-tastic is a platform for data amputation based on (Schouten et al., 2018). The authors propose a multivariate amputation procedure that simulates accurately missingness under various mechanisms (MCAR, MAR, MNAR), with precise control over which variables are affected, how much data is missing, and in what pattern. This method enables the building of missing data scenarios that closely reflect real-world conditions.

### 3.3 Missingness management

To deal with the issue of missing data management, three strategies may be considered:

- **Complete Case (CC):** Here no missing data is introduced. The covariates are complete and are kept that way.
- **Incomplete Case (IncC):** Also, missing data is introduced following one of Rubin’s missingness mechanisms at a time. The objective is to observe how missingness in the covariates affects the performance of the recoding methods under different target variables and distribution shifts.
- **Imputed Case (ImpC):** In this case, missing data in the covariates is imputed prior to recoding using MICE. MICE is chosen to estimate plausible values of the missing entries before variable recoding and analysis.

### 3.4 Simulation scenarios

The simulation scenarios involved aim to build databases A and B in order to investigate the performances of the three reconstruction algorithms regarding

1. Assumption on the distributions (AonD) of *Y*_*A*_ and *Y*_*B*_, with *S* indicating the same distribution and *D* indicating different distributions.
2. Sample size, denoted *n* = *n*_*A*_ + *n*_*B*_.
3. Ratio of observations in database *A* to those in database *B*, defined as *F* = *n*_*A*_*/n*_*B*_.
4. Strength of association between the covariates, controlled by *R*^2^.
5. Management of missing data (MMD) and the proportion of missingness in the covariates, denoted *P*.
6. Missing data mechanism (MDM): MCAR, MAR, or MNAR.

Table 2 fixes the values of the parameters of the different scenarios that were considered.

**Table 2.** Parameters of the simulation scenarios. MMD: Management of missing data, AonD: assumption on distribution,MDM: Missing Data Mechanism.

| Scen. | MMD | AonD | $n$ | $F$ | $R^2$ | $P$ | MDM |
| --- | --- | --- | --- | --- | --- | --- | --- |
| 1 | CC | S | 500 | $\{0.1, 1, 9\}$ | 0.9 | - | - |
| 2 | CC | S | $\{100, 200, 300, 400, 500, 1000\}$ | 1 | 0.9 | - | - |
| 3 | CC | S | 500 | 1 | $\{0.2, 0.5, 0.9\}$ | - | - |
| 4 | CC | D | $\{100, 200, 300, 400, 500, 1000\}$ | 1 | 0.8 | - | - |
| 5 | IncC | D | $\{100, 200, 300, 400, 500, 1000\}$ | 1 | 0.8 | $\{0.1, 0.5, 0.9\}$ | MCAR |
| 6 | IncC | D | $\{100, 200, 300, 400, 500, 1000\}$ | 1 | 0.8 | $\{0.1, 0.5, 0.9\}$ | MAR |
| 7 | IncC | D | $\{100, 200, 300, 400, 500, 1000\}$ | 1 | 0.8 | $\{0.1, 0.5, 0.9\}$ | MNAR |
| 8 | ImpC | D | $\{100, 200, 300, 400, 500, 1000\}$ | 1 | 0.8 | $\{0.1, 0.5, 0.9\}$ | MCAR |
| 9 | ImpC | D | $\{100, 200, 300, 400, 500, 1000\}$ | 1 | 0.8 | $\{0.1, 0.5, 0.9\}$ | MAR |
| 10 | ImpC | D | $\{100, 200, 300, 400, 500, 1000\}$ | 1 | 0.8 | $\{0.1, 0.5, 0.9\}$ | MNAR |

#### Remark 2

*The details on the tuning of algorithm parameters are found in Section 1 of the Supplementary Materials*.

### 3.5 Performance of recoding algorithms

Consider Database A and Database B a dataset generated following the steps of the previous section. If any, missing data are managed following a given one the strategies presented in Section 3.3: CC, IncC or ImpC. Once the missing data managed, the updated (by removing individuals or by imputing data) dataset looks like Sub-Figure 1b and the recoding of the database is possible using OTrecod, MICE, and kNN. These algorithms impute missing values in the outcome by values denoted (*y*_*A→B,i*_, *i* = *n*_*A*_ + 1, …, *n*_*A*_ + *n*_*B*_) for database *B* and (*y*_*B→A,i*_, *i* = 1, …, *n*_*A*_) for database *A*. The performance, in a given scenario, with a given management of the missing data and for a given algorithm, is assessed by:

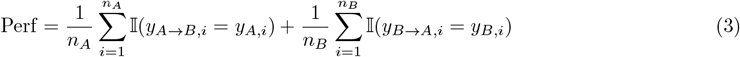

To specify the variability in the performances, the average among 30 simulations runs together with the standard deviation are computed.

## 4. Results

### 4.1 Complete Case (CC)

The results in Figure 2a reveal a clear dependence of recoding performance on the ratio F. For instance, when F is small (F=0.1), meaning dataset A has a tenth of the observations of dataset B, the OTrecod algorithm achieves the highest performance (0.702±0.039), indicating strong robustness to data imbalance even with limited samples. For the balanced case (F=1), OTrecod again outperforms MICE and kNN, suggesting that it remains the most reliable approach under symmetric sample sizes. However, when F becomes large, F=9, OTrecod’s performance declines, while MICE’s performance increases (0.646±0.015). kNN exhibits relatively stable but consistently lower performance across all scenarios.

**Figure 2.**
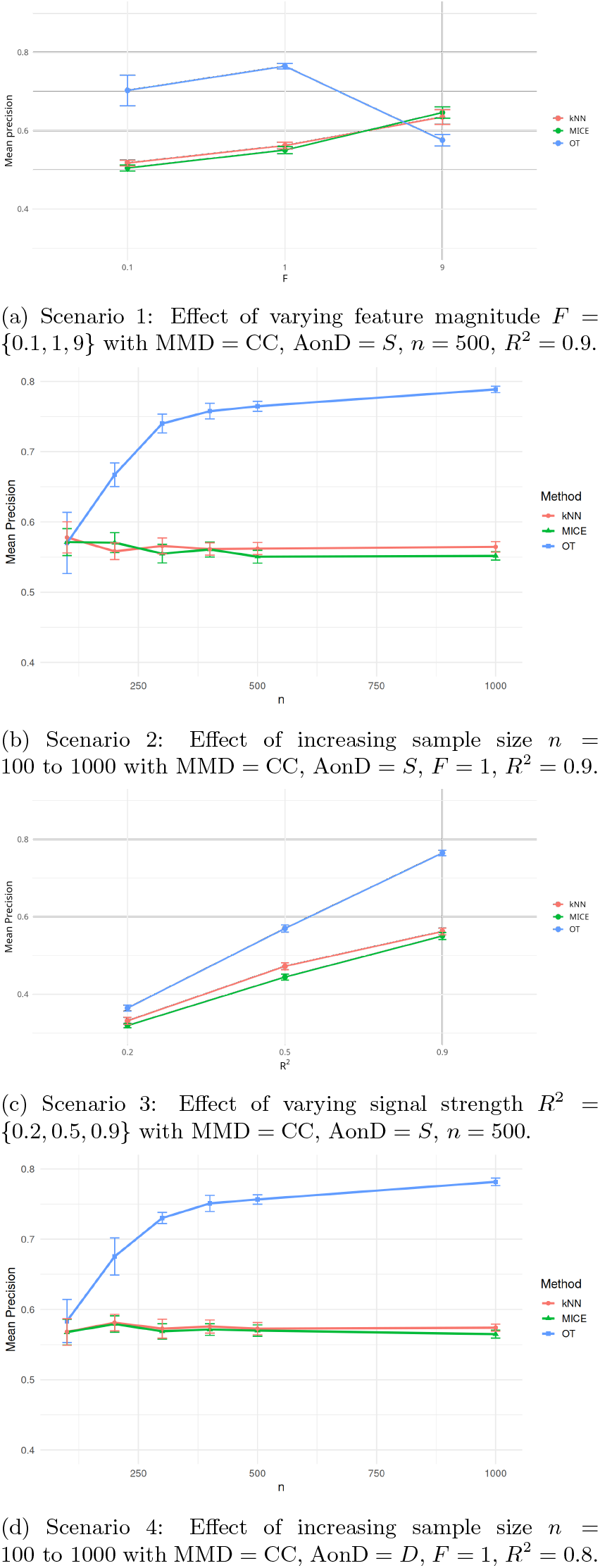
Scenarios 1–4 under complete cases (CC) with varying *n, F, R*^2^ and AonD

The results in Figure 2b demonstrate the evolution of recoding performance with increasing total sample size *n*. All three methods function similarly when *n* = 100, with kNN obtaining a slightly higher mean precision score (0.578 *±* 0.022). However, the OTrecod algorithm’s performance significantly improves as the total number of observations increases, outperforming both MICE and kNN from *n* = 200 onward. This improvement shows that OTrecod successfully captures relationships between variables and the underlying data structure in bigger datasets. MICE and kNN, on the other hand, exhibit comparatively steady but consistent performance as *n* increases, indicating limited scalability and sensitivity to sample size. These findings demonstrate that OTrecod has a considerable ability to gain from more data availability, attaining higher accuracy as the dataset grows, but MICE and kNN’s performance increases are limited.

Figure 2c presents the results that demonstrate how the coefficient of determination *R*^2^ affects data recoding. The OTrecod technique consistently beats both MICE and kNN across all levels of *R*^2^, but all methods exhibit improved results as *R*^2^ grows, demonstrating stronger correlations among covariates and outcome. In comparison to MICE and kNN, OTrecod achieves a much higher mean precision 0.3645 *±* 0.0078 when the variable connections are weak *R*^2^ = 0.2, indicating improved robustness under low-correlation situations. As *R*^2^ increases, this performance advantage becomes even more noticeable; at *R*^2^ = 0.9, OTrecod reaches 0.7646 *±* 0.0071, whereas MICE and kNN plateau at significantly lower values. These findings imply that as covariates-outcome correlations get stronger, OTrecod is better at identifying and taking advantage of the data’s underlying dependence structure, leading to higher recoding precision. Figure 2d serves as baseline for the comparison of the complete case with the different missingness scenarios of missingness in covariates.

### 4.2 Incomplete case (IncC)

When no imputation is done in the covariates before data recoding, the findings in Figures 3 show how various missingness techniques affect model performance. OTrecod consistently performs better than MICE and kNN under the MCAR condition for all *n* and *P*. OTrecod maintains comparatively consistent precision, demonstrating tolerance to random data loss, even if all approaches exhibit a little decrease in performance as *P* rises. These findings imply that whereas MICE and kNN show more sensitivity to the loss of observed information, OTrecod better maintains the underlying structure even when data are randomly missing.

**Figure 3.**
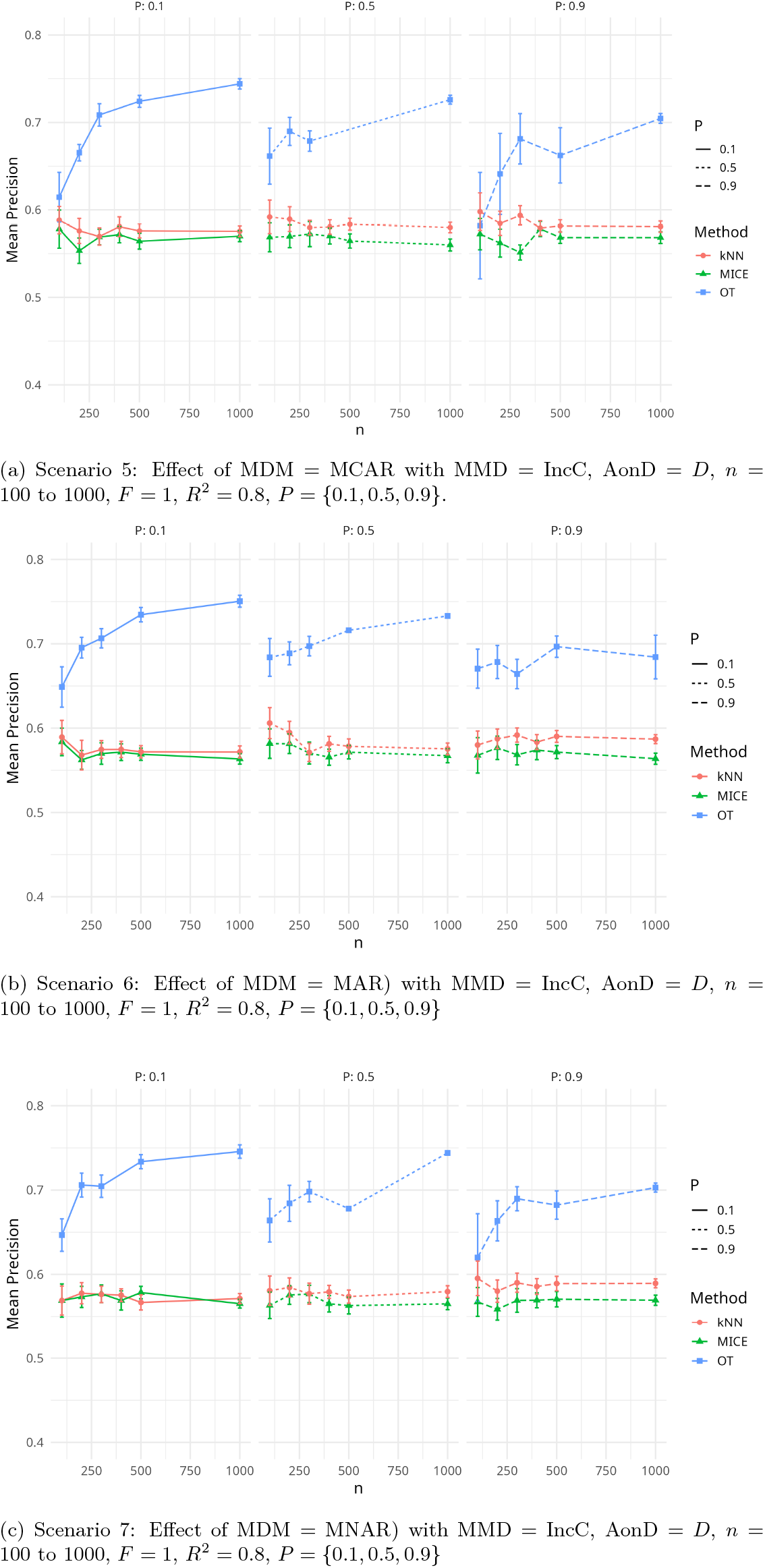
Scenarios 5–7 involve incomplete cases (IncC) under different MDM (MCAR, MAR, MNAR) with fixed *F, R*^2^, and varying proportions of missingness *P*.

Across all *P*s, OTrecod continues to produce better results in the MAR and MNAR settings, where *P* is dependent on observed or unobserved variables. As *P* increases, the disparity in performance between approaches becomes more noticeable, especially under MNAR, where unobserved connections worsen data distortion. In these situations, OTrecod still exhibits a significant level of resistance, but the lack of imputation clearly reduces accuracy for all approaches.

### 4.3 Imputed case(ImpC)

In the imputed scenarios, Figure 4, OTrecod generally achieves the highest performance across most *n* and *P*, particularly for moderate to larger *n*. kNN shows better results for smaller *n*. MICE and kNN show improved stability, with reduced variability across runs, though their absolute precision scores remain generally lower than OTrecod. The impact of imputation is most noticeable under high missingness, where it helps maintain more consistent results and mitigates drastic performance drops, especially for MICE and kNN.

**Figure 4.**
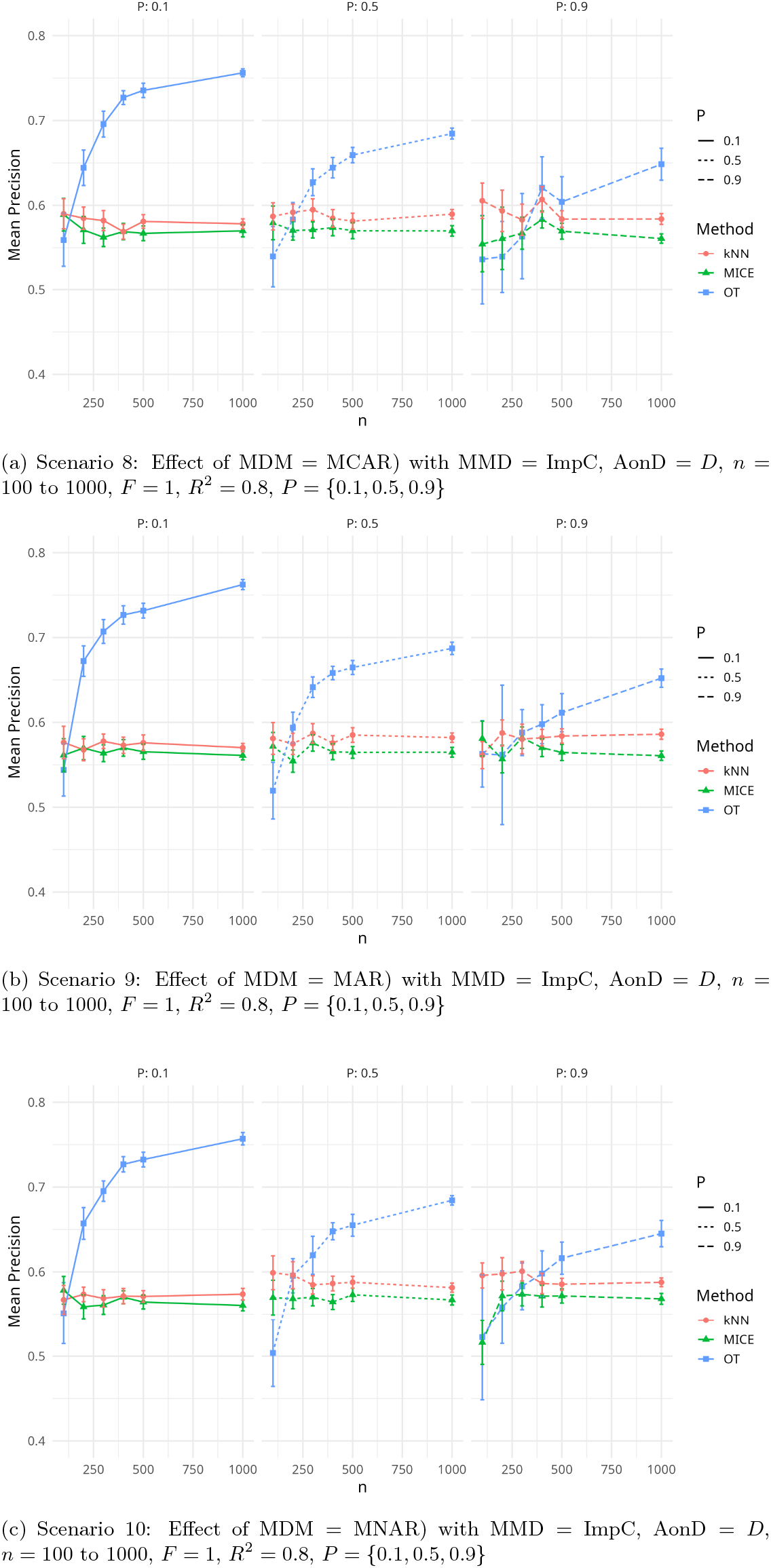
Scenarios 8–10 involve imputed cases (ImpC) under different MDM (MCAR, MAR, MNAR) with fixed *F, R*^2^, and varying proportions of missingness *P*.

## 5 Experimentation on real dataset

On real data, we do not have the true outcome and therefore we cannot compare recoding performance assessment. We are therefore interested in the agreement of the results obtained by the three approaches according to two criteria: Wasserstein distances to assess the pairwise agreement in terms of distribution and, at an individual level, the percentage of overlaps between two or three approaches.

### 5.1 Description of the NCDS dataset

The experimentation is conducted using data from the National Child Development Study (NCDS), which is a long-term longitudinal survey that has followed the lives of over 17,000 individuals born in a single week of 1958 across England, Scotland, and Wales. The NCDS collects extensive information on multiple life domains, including physical and educational development, socioeconomic circumstances, employment trajectories, family composition, health status, well-being, social participation, and attitudes. Among the numerous variables recorded, particular attention is given to the participants’ social class, measured using two occupationally based classification systems collected at each wave of data gathering.

The first of these was the Goldthorp social class ‘90 scale (GSS90), an ordered scale with 11 categories, while the second one was the RGs social Class ‘91 scale (RGS91), an ordered scale with only 6 categories. The initial dataset was randomly divided in two datasets of the same size. Social class of subjects was considered as the outcome only supposed measured with the GSS90 scale in the first dataset generated (DB1), while it was assumed to be only supposed measured in the RGS91 scale for the second one (DB2).

The aim of the study was here to solve the recoding problem in each of the two datasets in order to obtain a unique scale in both bases. DB1 contains 4 covariates and 5476 observations while DB2 contains 4 covariates and 365 observations.

### 5.2 Results

In the complete case (CC) scenario, only observations with fully recorded values for all covariates and the social class variable were retained for analysis. *Y*_*A*_ contains 307 observations while *Y*_*B*_, 4013 observations. Concordance between the recoding methods varies across *Y*_*A*_ and *Y*_*B*_ (Table 3). For *Y*_*A*_, OTrecod and kNN agree on 26.40% of observations, OTrecod and MICE on 35.50%, and kNN and MICE on 20.50%. Only 10.10% of observations are classified identically by all three methods, while 37.80% are uniquely classified by a single method. For *Y*_*B*_, OTrecod and kNN agree on 35.10%, OTrecod and MICE on 39.30%, and kNN and MICE on 25.40%, with 14.00% classified identically by all three methods and 28.20% classified uniquely by a single method. These values indicate that, in the complete case, full agreement across all three methods is limited, while pairwise agreement is moderate.

**Table 3.** Concordance percentages between OTrecod, kNN, and MICE for *Y*_*A*_ and *Y*_*B*_.

| Cases | OT and kNN | | OT and MICE | | kNN and MICE | | OT, kNN and MICE | | $n$ | |
| --- | --- | --- | --- | --- | --- | --- | --- | --- | --- | --- |
| | $Y_A$ | $Y_B$ | $Y_A$ | $Y_B$ | $Y_A$ | $Y_B$ | $Y_A$ | $Y_B$ | $Y_A$ | $Y_B$ |
| CC | 26.40 | 35.10 | 35.50 | 39.30 | 20.50 | 25.40 | 10.10 | 14.00 | 307 | 4013 |
| IncC | 27.90 | 39.50 | 33.20 | 41.20 | 17.80 | 31.30 | 9.30 | 18.40 | 365 | 4670 |
| ImpC | 26.80 | 39.50 | 28.20 | 37.80 | 18.10 | 32.90 | 8.80 | 18.60 | 365 | 4670 |

The incomplete case (IncC) scenario considers the original dataset containing missing values in covariates. In this configuration, incomplete observations are preserved. A total of 5035 observations for 6 variables are recorded, of which *Y*_*A*_, 365 observations and *Y*_*B*_, 4670 observations. In the incomplete case (IncC) scenario, where missing data are present, the patterns of concordance are similar but slightly higher for *Y*_*B*_ (Table 3). For *Y*_*A*_, OTrecod and kNN agree on 27.90% of observations, OTrecod and MICE on 33.20%, and kNN and MICE on 17.80%, with 9.30% of observations classified identically by all three methods and 39.70% uniquely classified by a single method. For *Y*_*B*_, OTrecod and kNN agree on 39.50%, OTrecod and MICE on 41.20%, and kNN and MICE on 31.30%, with 18.40% classified identically by all three methods and 24.80% uniquely classified. This shows that incomplete data slightly increases pairwise and triple agreement for *Y*_*B*_ compared to *Y*_*A*_.

In the imputed case (ImpC) scenario, missing data in the covariates were estimated using MICE before recoding the social class variable. The resulting imputed covariates were then used to solve the recoding problem in the target variables. A total of 5,035 observations for 6 variables are recorded of which *Y*_*A*_, 365 observations and *Y*_*B*_, 4670 observations. In the imputed case (ImpC) scenario, using MICE to impute covariates, concordance percentages remain comparable to the incomplete case (Table 3). For *Y*_*A*_, OTrecod and kNN agree on 26.80% of observations, OTrecod and MICE on 28.20%, and kNN and MICE on 18.10%, with 8.80% classified identically by all three methods and 44.40% uniquely classified. For *Y*_*B*_, OTrecod and kNN agree on 39.50%, OTrecod and MICE on 37.80%, and kNN and MICE on 32.90%, with 18.60% of observations classified identically by all three methods and 27.00% classified uniquely. These percentages indicate that imputation reduces the differences between OTrecod and MICE for *Y*_*A*_, but overall full concordance remains limited.

Globally, whatever the method and the scenario, results are less consistent when considering recoding *Y*_*A*_. This makes sense because of the lower number of observations used to learn and the higher number of observations to be recoded. Imputing the missing values for the covariates does not impact so much the results. The highest agreements are observed for OTrecod and kNN, and for OTrecod and MICE. Regarding OTrecod and kNN, this is consistent with the fact that OTrecod partly relies on kNN. More surprisingly, OTrecod and MICE also obtains concordant results. On the oher hand, MICE and kNN disagree more frequently. It appears that OTrecod seems to take benefit from both MICE and kNN in a synergistic way.

As shown in Table 4, the Wasserstein distances reveal clear differences in how the three methods compare across variables and scenarios. For *Y*_*B*_, the largest discrepancies are observed between kNN and the two other methods, with distances between 0.39 (OT-kNN, ImpC) and 0.57 (OT-kNN, IncC). In contrast, OTrecod and MICE exhibit the smallest distances (around 0.02) for *Y*_*A*_ across all scenarios, indicating that these two methods generate the most similar distributions for this variable. The results suggest that while all methods behave similarly for *Y*_*A*_, the distributions for *Y*_*B*_ diverge more substantially, especially considering kNN.

**Table 4.**
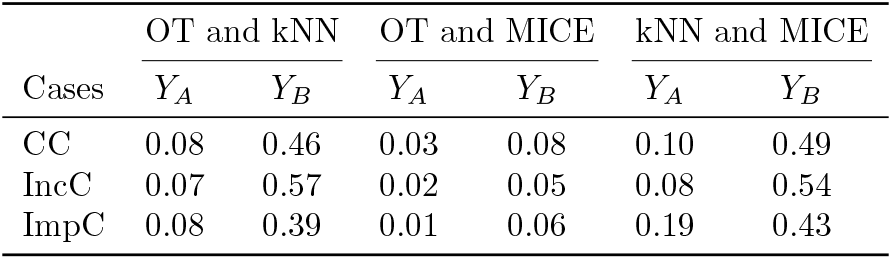
Wasserstein distances between methods for different cases.

Results in terms of distribution are consistent with the previous results at individual level regarding the impact of imputation of missing values in covariates and the difficulty to recode *Y*_*A*_. However, we can note that the distribution obtained by OTrecod is not so close from the one obtained by kNN.

## 6 Discussion and Conclusion

Variable recoding is an essential problem in data merging. Among different approaches, the choice between statistically-oriented methods such as optimal transport and machine learning methods in this context is one addressed here. The choice of method to be used in different scenarios is an essential question. The study investigated the performance of three methods, OTrecod, MICE and kNN, in the recoding problem under varying conditions, including the group size ratio (*F*), total sample size (*n*), and the determination coefficient (*R*^2^), as well as incomplete and imputed cases. OTrecod consistently outperforms the other methods across most conditions due to its ability to exploit distributional structure.

Varying the group size ratio had a notable impact on methods performance. More precisely, OTrecod excels when the two group sizes are balanced. Increasing the total sample size improved performance for all methods. OTrecod showed the most substantial gains with larger datasets, while MICE and kNN experienced more modest improvements. Higher values for the determination coefficient (*R*^2^), representing stronger covariates-outcome relationships, generally increased performance for all methods. The missingness mechanisms (MCAR, MAR, MNAR) in data have a strong influence on the performance of the methods. The results suggest that imputation does not necessarily increase performance, particularly for OTrecod, which already handles partial data structures effectively. In some cases, using incomplete data directly (without imputation) preserved more of the true distributional information that OTrecod exploits. However, imputation improved stability and reduced variability across methods and runs, making results more predictable, especially for MICE and kNN, which rely on complete-case analysis. Considering the NCDS dataset, the imputation strategy does not influence the agreement between methods. Moreover, the results obtained with OTrecod are the most consistent with the other two methods, whereas kNN and MICE seem to disagree more often.

In light of all these results, we strongly recommend to use OTrecod after imputing missing values in the covariates. We believe that this work offers an opportunity to reconcile heterogeneous datasets in terms of coding qualitative information and thus contribute to putting FAIR principles, mainly Interoperability and Re-usability, into practice.

## Author Contributions

Flore N’kam Suguem (Conceptualization [lead], Software [lead], Formal analysis [lead], Investigation [lead], Data curation [lead], Visualization [lead], Writing—original draft [lead], Writing—review, editing [equal]); Sébastien Déjean (Supervision [lead], Funding acquisition [equal], Writing—introduction [equal], Writing—review, editing [equal]); Philippe Saint Pierre (Methodology [lead], Funding acquisition [equal]), Writing—review,editing [equal]; and Nicolas Savy (Conceptualization [equal], Methodology [equal], Supervision [lead], Funding acquisition [equal], Writing—review,editing [equal], Paper structure and template design [equal]).

## Data Availability

The National Child Development Study dataset is included in the R package OTrecod from the Comprehensive R Archive Network (CRAN): https://cran.r-project.org.

## Conflict of interest

The authors declare that they have no competing interests.

## Grant Information

This work was supported by the FAIROMICS project, funded by the European Union’s Horizon 2020 Research and Innovation program under the Marie Sklodowska-Curie Grant Agreement No. 101120449.

## Notes

### Competing Interest Statement

The authors have declared no competing interest.

