## Supplementary materials for "Comparing optimal transport and machine learning approaches for databases merging in scenarios involving missing data in covariates. Application to Medical Research"

### Comparing Optimal Transport and Machine Learning Approaches for Data Merging Under Missing Covariate Scenarios: Application to Medical Research
### Supplementary Materials

Flore N'kam Suguem [1,2,4,*], Sébastien Déjean [1,2], Philippe Saint Pierre [1,2], and Nicolas Savy [1,3 *]

[1]Institut de Mathématiques de Toulouse, UMR521, Université de Toulouse, CNRS, France
[2]Université de Toulouse UT, F-31062 Toulouse Cedex 9, France
[3]Université Toulouse Jean-Jaurès UT2J, F-31058 Toulouse, France
[4]Department of Physics and Astronomy, University of Bologna, 40127 Bologna, Italy

January 23, 2026

### 1 Code and Packages

The following packages are used to get the results described in this manuscript: **Software:** All analyses were conducted in R (version 4.4.1) using the following packages:

- OTrecod (version 0.1.2)
- mice (version 3.18.0)
- VIM (version 6.2.2)

All packages are available from the Comprehensive R Archive Network (CRAN): `https://cran.r-project.org`.

#### OTrecod

- **Type** = R-JOINT
- **Distance** = Hamming
- **Percent closest k-NN** = 100%
- **Relaxation parameter** = YES
- **Relaxation value** = 0.1 (set according to the distribution of our dataset)
- **Regularization term** = 0
- **Aggregation tol cov** = 0.3
- **DB imputed** = BOTH

#### MICE 3.0

In the `mice` function, the following parameters are specified:

- `method = 'rf'`: Specifies that random forest should be used for imputation.
- `m = 5`: Specifies that 5 imputed datasets should be created.
- `maxit = 5`: Specifies that 5 iterations should be performed.

#### VIM

- **k**: set at 5
- **distVar**: The distance metric typically Euclidean.
- **impVar**: The set of variables to use for imputation (FALSE).

The other parameters were left unchanged as originally set in the R-package VIM. A smaller value of k may overfit, while a larger k might smooth out the imputation.

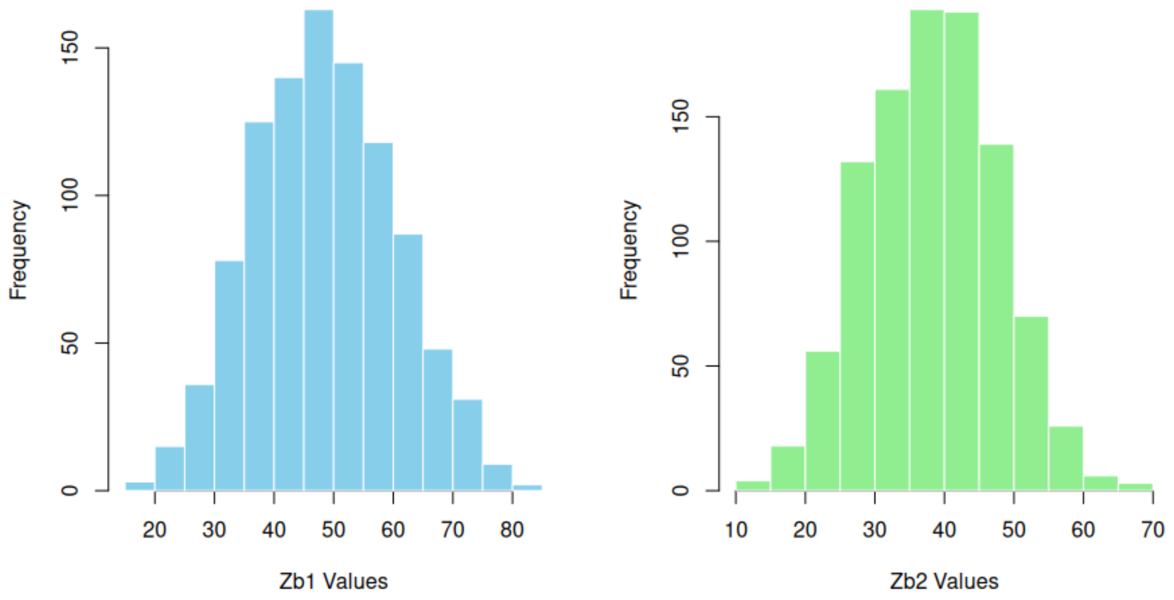


(a) Histogram of the values before categorisation

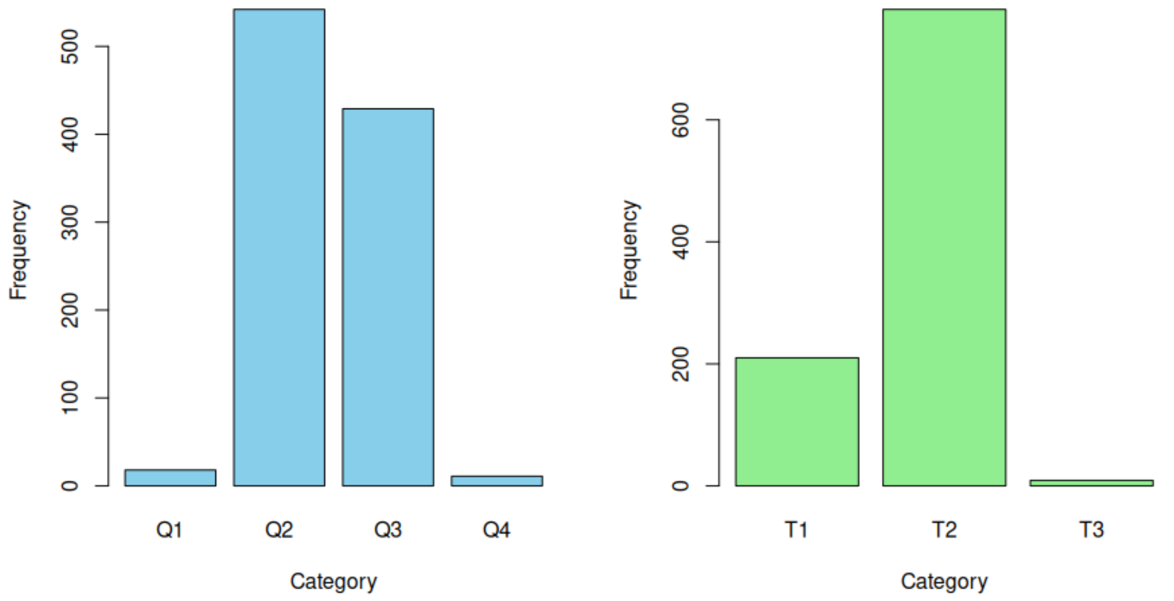


(b) Histogram of the categories

Figure 1: Histograms of the distribution of the outcome Zb1 and Zb2 for $n = 1000$

### 2 Tables of Results

Table 1: Scenario 1: Effect of varying feature magnitude $F = \{0.1, 1, 9\}$ with MMD = CC, AonD = $S$, $n = 500$, $R^2 = 0.9$, see Figure 2(a) of the article.

| F | OT (mean ± error) | MICE (mean ± error) | kNN (mean ± error) |
|---|---|---|---|
| 0.1 | **0.7028 ± 0.0390** | 0.5049 ± 0.0077 | 0.5175 ± 0.0071 |
| 1 | **0.7646 ± 0.0071** | 0.5505 ± 0.0091 | 0.5621 ± 0.0086 |
| 9 | 0.5759 ± 0.0146 | **0.6464 ± 0.0146** | 0.6349 ± 0.0187 |

Table 2: Scenario 2: Effect of increasing sample size $n = 100$ to 1000 with MMD = CC, AonD = $S$, $F = 1$, $R^2 = 0.9$, see Figure 2(b) of the article.

| I | OT (mean ± error) | MICE (mean ± error) | kNN (mean ± error) |
|---|---|---|---|
| 100 | 0.5700 ± 0.0435 | 0.5713 ± 0.0191 | **0.5780 ± 0.0222** |
| 200 | **0.6671 ± 0.0169** | 0.5705 ± 0.0142 | 0.5582 ± 0.0118 |
| 300 | **0.7400 ± 0.0133** | 0.5549 ± 0.0132 | 0.5658 ± 0.0113 |
| 400 | **0.7576 ± 0.0111** | 0.5608 ± 0.0107 | 0.5614 ± 0.0087 |
| 500 | **0.7646 ± 0.0071** | 0.5505 ± 0.0091 | 0.5621 ± 0.0086 |
| 1000 | **0.7886 ± 0.0046** | 0.5517 ± 0.0060 | 0.5645 ± 0.0075 |

Table 3: Scenario 3: Effect of varying signal strength $R^2 = \{0.2, 0.5, 0.9\}$ with MMD = CC, AonD = $S$, $n = 500$, see Figure 2(c) of the article.

| $R^2$ | OT (mean ± error) | MICE (mean ± error) | kNN (mean ± error) |
|---|---|---|---|
| 0.2 | **0.3645 ± 0.0078** | 0.3194 ± 0.0058 | 0.3315 ± 0.00896 |
| 0.5 | **0.5699 ± 0.0092** | 0.4445 ± 0.0078 | 0.4724 ± 0.00901 |
| 0.9 | **0.7646 ± 0.0071** | 0.5505 ± 0.0091 | 0.5621 ± 0.00860 |

Table 4: Scenario 4: Effect of increasing sample size $n = 100$ to 1000 with MMD = CC, AonD = $D$, $F = 1$, $R^2 = 0.8$, see Figure 2(d) of the article.

| $n$ | OT | MICE | kNN |
|---|---|---|---|
| 100 | **0.5835 ± 0.0306** | 0.5677 ± 0.0182 | 0.5680 ± 0.0190 |
| 200 | **0.6753 ± 0.0264** | 0.5792 ± 0.0116 | 0.5812 ± 0.0118 |
| 300 | **0.7301 ± 0.0079** | 0.5689 ± 0.0109 | 0.5727 ± 0.0135 |
| 400 | **0.7509 ± 0.0115** | 0.5715 ± 0.0083 | 0.5757 ± 0.0093 |
| 500 | **0.7566 ± 0.0067** | 0.5699 ± 0.0081 | 0.5727 ± 0.0087 |
| 1000 | **0.7816 ± 0.0054** | 0.5649 ± 0.0056 | 0.5741 ± 0.0051 |

Table 5: Scenario 5: Effect of MDM = MCAR with MMD = IncC, AonD = $D$, $n$ = 100 to 1000, $F = 1$, $R^2 = 0.8$, $P = \{0.1, 0.5, 0.9\}$, see Figure 3(a) of the article.

| $P$ | $n$ | Method | | |
|---|---|---|---|---|
| | | **OT** | **MICE** | **kNN** |
| 10% | 100 | $\mathbf{0.6145 \pm 0.0284}$ | $0.5780 \pm 0.0218$ | $0.5883 \pm 0.0156$ |
| | 200 | $\mathbf{0.6654 \pm 0.0094}$ | $0.5533 \pm 0.0145$ | $0.5760 \pm 0.0144$ |
| | 300 | $\mathbf{0.7086 \pm 0.0128}$ | $0.5689 \pm 0.0088$ | $0.5696 \pm 0.0099$ |
| | 400 | $\mathbf{0.7194 \pm 0.0113}$ | $0.5718 \pm 0.0094$ | $0.5808 \pm 0.0114$ |
| | 500 | $\mathbf{0.7242 \pm 0.0068}$ | $0.5641 \pm 0.0089$ | $0.5759 \pm 0.0079$ |
| | 1000 | $\mathbf{0.7441 \pm 0.0060}$ | $0.5699 \pm 0.0062$ | $0.5755 \pm 0.0062$ |
| 50% | 100 | $\mathbf{0.6615 \pm 0.0320}$ | $0.5687 \pm 0.0166$ | $0.5920 \pm 0.0192$ |
| | 200 | $\mathbf{0.6898 \pm 0.0160}$ | $0.5697 \pm 0.0129$ | $0.5895 \pm 0.0142$ |
| | 300 | $\mathbf{0.6787 \pm 0.0117}$ | $0.5723 \pm 0.0143$ | $0.5798 \pm 0.0084$ |
| | 400 | $\mathbf{0.7525 \pm 0.0110}$ | $0.5804 \pm 0.0083$ | $0.5808 \pm 0.0114$ |
| | 500 | $\mathbf{0.7000 \pm 0.0073}$ | $0.5645 \pm 0.0081$ | $0.5837 \pm 0.0069$ |
| | 1000 | $\mathbf{0.7260 \pm 0.0050}$ | $0.5599 \pm 0.0068$ | $0.5799 \pm 0.0061$ |
| 90% | 100 | $\mathbf{0.5820 \pm 0.0609}$ | $0.5723 \pm 0.0180$ | $0.5980 \pm 0.0215$ |
| | 200 | $\mathbf{0.6413 \pm 0.0463}$ | $0.5620 \pm 0.0160$ | $0.5847 \pm 0.0138$ |
| | 300 | $\mathbf{0.6813 \pm 0.0289}$ | $0.5513 \pm 0.0086$ | $0.5938 \pm 0.0110$ |
| | 400 | $\mathbf{0.6782 \pm 0.0267}$ | $0.5782 \pm 0.0087$ | $0.5793 \pm 0.0085$ |
| | 500 | $\mathbf{0.6623 \pm 0.0316}$ | $0.5683 \pm 0.0066$ | $0.5817 \pm 0.0071$ |
| | 1000 | $\mathbf{0.7046 \pm 0.0056}$ | $0.5683 \pm 0.0066$ | $0.5809 \pm 0.0065$ |

Table 6: Scenario 6: Effect of MDM = MAR) with MMD = IncC, AonD = $D$, $n$ = 100 to 1000, $F$ = 1, $R^2 = 0.8$, $P = \{0.1, 0.5, 0.9\}$, see Figure 3(b) of the article.

| $P$ | $n$ | Method | | |
|---|---|---|---|---|
| | | OT | MICE | kNN |
| 10% | 100 | $\mathbf{0.6488 \pm 0.0239}$ | $0.5837 \pm 0.0163$ | $0.5893 \pm 0.0198$ |
| | 200 | $\mathbf{0.6954 \pm 0.0123}$ | $0.5623 \pm 0.0111$ | $0.5680 \pm 0.0175$ |
| | 300 | $\mathbf{0.7065 \pm 0.0114}$ | $0.5699 \pm 0.0127$ | $0.5747 \pm 0.0105$ |
| | 400 | $\mathbf{0.7236 \pm 0.0085}$ | $0.5715 \pm 0.0099$ | $0.5748 \pm 0.0096$ |
| | 500 | $\mathbf{0.7345 \pm 0.0086}$ | $0.5691 \pm 0.0072$ | $0.5719 \pm 0.0073$ |
| | 1000 | $\mathbf{0.7505 \pm 0.0071}$ | $0.5635 \pm 0.0061$ | $0.5717 \pm 0.0071$ |
| 50% | 100 | $\mathbf{0.6839 \pm 0.0224}$ | $0.5817 \pm 0.0175$ | $0.6060 \pm 0.0184$ |
| | 200 | $\mathbf{0.6887 \pm 0.0136}$ | $0.5815 \pm 0.0116$ | $0.5947 \pm 0.0134$ |
| | 300 | $\mathbf{0.6973 \pm 0.0116}$ | $0.5704 \pm 0.0129$ | $0.5712 \pm 0.0107$ |
| | 400 | $\mathbf{0.7270 \pm 0.0090}$ | $0.5656 \pm 0.0096$ | $0.5813 \pm 0.0089$ |
| | 500 | $\mathbf{0.7160 \pm 0.0925}$ | $0.5715 \pm 0.0079$ | $0.5785 \pm 0.0087$ |
| | 1000 | $\mathbf{0.7330 \pm 0.0070}$ | $0.5673 \pm 0.0083$ | $0.5754 \pm 0.0069$ |
| 90% | 100 | $\mathbf{0.6705 \pm 0.0232}$ | $0.5677 \pm 0.0210$ | $0.5800 \pm 0.0165$ |
| | 200 | $\mathbf{0.6784 \pm 0.0197}$ | $0.5767 \pm 0.0139$ | $0.5873 \pm 0.0117$ |
| | 300 | $\mathbf{0.6642 \pm 0.0174}$ | $0.5684 \pm 0.0120$ | $0.5918 \pm 0.0084$ |
| | 400 | $\mathbf{0.6827 \pm 0.0131}$ | $0.5740 \pm 0.0114$ | $0.5835 \pm 0.0089$ |
| | 500 | $\mathbf{0.6966 \pm 0.0127}$ | $0.5716 \pm 0.0077$ | $0.5902 \pm 0.0070$ |
| | 1000 | $\mathbf{0.6843 \pm 0.0258}$ | $0.5638 \pm 0.0066$ | $0.5870 \pm 0.0053$ |

Table 7: Scenario 7: Effect of MDM = MNAR) with MMD = IncC, AonD = $D$, $n$ = 100 to 1000, $F$ = 1, $R^2 = 0.8$, $P = \{0.1, 0.5, 0.9\}$, see Figure 3(c) of the article.

| $P$ | $n$ | Method | | |
|---|---|---|---|---|
| | | OT | MICE | kNN |
| 10% | 100 | $\mathbf{0.6467 \pm 0.0193}$ | $0.5690 \pm 0.0197$ | $0.5690 \pm 0.0173$ |
| | 200 | $\mathbf{0.7059 \pm 0.0142}$ | $0.5733 \pm 0.0125$ | $0.5777 \pm 0.0125$ |
| | 300 | $\mathbf{0.7045 \pm 0.0132}$ | $0.5770 \pm 0.0105$ | $0.5763 \pm 0.0097$ |
| | 400 | $\mathbf{0.7257 \pm 0.0105}$ | $0.5691 \pm 0.0113$ | $0.5753 \pm 0.0076$ |
| | 500 | $\mathbf{0.7336 \pm 0.0083}$ | $0.5785 \pm 0.0074$ | $0.5667 \pm 0.0089$ |
| | 1000 | $\mathbf{0.7457 \pm 0.0079}$ | $0.5653 \pm 0.0052$ | $0.5713 \pm 0.0059$ |
| 50% | 100 | $\mathbf{0.6640 \pm 0.0257}$ | $0.5633 \pm 0.0156$ | $0.5807 \pm 0.0173$ |
| | 200 | $\mathbf{0.6843 \pm 0.0213}$ | $0.5755 \pm 0.0110$ | $0.5845 \pm 0.0114$ |
| | 300 | $\mathbf{0.6981 \pm 0.0121}$ | $0.5768 \pm 0.0102$ | $0.5771 \pm 0.0125$ |
| | 400 | $\mathbf{0.6925 \pm 0.0343}$ | $0.5652 \pm 0.0097$ | $0.5793 \pm 0.0077$ |
| | 500 | $\mathbf{0.6780 \pm 0.03}$ | $0.5630 \pm 0.0099$ | $0.5737 \pm 0.0078$ |
| | 1000 | $\mathbf{0.7440 \pm 0.03}$ | $0.5651 \pm 0.0069$ | $0.5796 \pm 0.0069$ |
| 90% | 100 | $\mathbf{0.6200 \pm 0.0519}$ | $0.5673 \pm 0.0171$ | $0.5953 \pm 0.0204$ |
| | 200 | $\mathbf{0.6635 \pm 0.0238}$ | $0.5587 \pm 0.0129$ | $0.5802 \pm 0.0132$ |
| | 300 | $\mathbf{0.6897 \pm 0.0142}$ | $0.5691 \pm 0.0140$ | $0.5901 \pm 0.0113$ |
| | 400 | $\mathbf{0.6985 \pm 0.0103}$ | $0.5693 \pm 0.0089$ | $0.5857 \pm 0.0093$ |
| | 500 | $\mathbf{0.6822 \pm 0.0168}$ | $0.5706 \pm 0.0091$ | $0.5891 \pm 0.0086$ |
| | 1000 | $\mathbf{0.7029 \pm 0.0053}$ | $0.5693 \pm 0.0060$ | $0.5894 \pm 0.0052$ |

Table 8: Scenario 8: Effect of MDM = MCAR) with MMD = ImpC, AonD = $D$, $n$ = 100 to 1000, $F$ = 1, $R^2 = 0.8$, $P = \{0.1, 0.5, 0.9\}$, see Figure 4(a) of the article.

| $P$ | $n$ | Method | | |
|---|---|---|---|---|
| | | OT | MICE | kNN |
| 10% | 100 | $0.5591 \pm 0.0317$ | $0.5890 \pm 0.0192$ | $\mathbf{0.5900 \pm 0.0177}$ |
| | 200 | $\mathbf{0.6443 \pm 0.0209}$ | $0.5710 \pm 0.0158$ | $0.5848 \pm 0.0131$ |
| | 300 | $\mathbf{0.6958 \pm 0.0152}$ | $0.5623 \pm 0.0109$ | $0.5821 \pm 0.0118$ |
| | 400 | $\mathbf{0.7270 \pm 0.0081}$ | $0.5691 \pm 0.0096$ | $0.5690 \pm 0.0088$ |
| | 500 | $\mathbf{0.7355 \pm 0.0084}$ | $0.5670 \pm 0.0086$ | $0.5810 \pm 0.0080$ |
| | 1000 | $\mathbf{0.7562 \pm 0.0047}$ | $0.5699 \pm 0.0072$ | $0.5783 \pm 0.0058$ |
| 50% | 100 | $0.5391 \pm 0.0359$ | $0.5793 \pm 0.0199$ | $\mathbf{0.5870 \pm 0.0160}$ |
| | 200 | $0.5833 \pm 0.0201$ | $0.5703 \pm 0.0112$ | $\mathbf{0.5918 \pm 0.0098}$ |
| | 300 | $\mathbf{0.6958 \pm 0.0152}$ | $0.5710 \pm 0.0095$ | $0.5949 \pm 0.0127$ |
| | 400 | $\mathbf{0.6444 \pm 0.0121}$ | $0.5738 \pm 0.0097$ | $0.5847 \pm 0.0103$ |
| | 500 | $\mathbf{0.6592 \pm 0.0090}$ | $0.5701 \pm 0.0093$ | $0.5813 \pm 0.0094$ |
| | 1000 | $\mathbf{0.6846 \pm 0.0064}$ | $0.5699 \pm 0.0061$ | $0.5896 \pm 0.0054$ |
| 90% | 100 | $0.5356 \pm 0.0526$ | $0.5543 \pm 0.0334$ | $\mathbf{0.6055 \pm 0.0209}$ |
| | 200 | $0.5388 \pm 0.0422$ | $0.5607 \pm 0.0371$ | $\mathbf{0.5935 \pm 0.0244}$ |
| | 300 | $0.5633 \pm 0.0506$ | $0.5672 \pm 0.0190$ | $\mathbf{0.5830 \pm 0.0185}$ |
| | 400 | $\mathbf{0.6208 \pm 0.0364}$ | $0.5833 \pm 0.0100$ | $0.6069 \pm 0.0150$ |
| | 500 | $\mathbf{0.6041 \pm 0.0297}$ | $0.5695 \pm 0.0093$ | $0.5838 \pm 0.0100$ |
| | 1000 | $\mathbf{0.6485 \pm 0.0188}$ | $0.5609 \pm 0.0055$ | $0.5839 \pm 0.0063$ |

Table 9: Scenario 9: Effect of MDM = MAR) with MMD = ImpC, AonD = $D$, $n$ = 100 to 1000, $F$ = 1, $R^2 = 0.8$, $P = \{0.1, 0.5, 0.9\}$, see Figure 4(b) of the article.

| $P$ | $n$ | Method | | |
|---|---|---|---|---|
| | | OT | MICE | kNN |
| 10% | 100 | $0.5440 \pm 0.0309$ | $0.5613 \pm 0.0195$ | $\mathbf{0.5763 \pm 0.0191}$ |
| | 200 | $\mathbf{0.6722 \pm 0.0180}$ | $0.5697 \pm 0.0137$ | $0.5677 \pm 0.0129$ |
| | 300 | $\mathbf{0.7071 \pm 0.0140}$ | $0.5636 \pm 0.0099$ | $0.5777 \pm 0.0086$ |
| | 400 | $\mathbf{0.7266 \pm 0.0108}$ | $0.5699 \pm 0.0098$ | $0.5730 \pm 0.0096$ |
| | 500 | $\mathbf{0.7316 \pm 0.0087}$ | $0.5654 \pm 0.0090$ | $0.5759 \pm 0.0092$ |
| | 1000 | $\mathbf{0.7624 \pm 0.0060}$ | $0.5610 \pm 0.0051$ | $0.5702 \pm 0.0050$ |
| 50% | 100 | $0.5195 \pm 0.0334$ | $0.5717 \pm 0.0164$ | $\mathbf{0.5810 \pm 0.0187}$ |
| | 200 | $\mathbf{0.5937 \pm 0.0183}$ | $0.5543 \pm 0.0131$ | $0.5747 \pm 0.0125$ |
| | 300 | $\mathbf{0.6415 \pm 0.0121}$ | $0.5756 \pm 0.0096$ | $0.5870 \pm 0.0115$ |
| | 400 | $\mathbf{0.6582 \pm 0.0079}$ | $0.5654 \pm 0.0093$ | $0.5756 \pm 0.0087$ |
| | 500 | $\mathbf{0.6647 \pm 0.0082}$ | $0.5646 \pm 0.0069$ | $0.5851 \pm 0.0086$ |
| | 1000 | $\mathbf{0.6872 \pm 0.0073}$ | $0.5648 \pm 0.0057$ | $0.5820 \pm 0.0055$ |
| 90% | 100 | $0.5629 \pm 0.0391$ | $\mathbf{0.5809 \pm 0.0205}$ | $0.5613 \pm 0.0159$ |
| | 200 | $0.5617 \pm 0.0822$ | $0.5567 \pm 0.0163$ | $\mathbf{0.5875 \pm 0.0153}$ |
| | 300 | $\mathbf{0.5880 \pm 0.0271}$ | $0.5819 \pm 0.0127$ | $0.5803 \pm 0.0175$ |
| | 400 | $\mathbf{0.5979 \pm 0.0229}$ | $0.5701 \pm 0.0105$ | $0.5819 \pm 0.0097$ |
| | 500 | $\mathbf{0.6114 \pm 0.0225}$ | $0.5645 \pm 0.0095$ | $0.5839 \pm 0.0085$ |
| | 1000 | $\mathbf{0.6521 \pm 0.0108}$ | $0.5607 \pm 0.0055$ | $0.5860 \pm 0.0059$ |

Table 10: Scenario 10: Effect of MDM = MNAR) with MMD = ImpC, AonD = $D$, $n$ = 100 to 1000, $F = 1$, $R^2 = 0.8$, $P = \{0.1, 0.5, 0.9\}$, see Figure 4(c) of the article.

| $P$ | $n$ | Method | | |
|---|---|---|---|---|
| | | OT | MICE | kNN |
| 10% | 100 | $0.5511 \pm 0.0362$ | $0.5783 \pm 0.0163$ | $\mathbf{0.5670 \pm 0.0171}$ |
| | 200 | $\mathbf{0.6572 \pm 0.0187}$ | $0.5588 \pm 0.0141$ | $0.5735 \pm 0.0085$ |
| | 300 | $\mathbf{0.6952 \pm 0.0120}$ | $0.5610 \pm 0.0111$ | $0.5688 \pm 0.0103$ |
| | 400 | $\mathbf{0.7268 \pm 0.0090}$ | $0.5702 \pm 0.0077$ | $0.5715 \pm 0.0089$ |
| | 500 | $\mathbf{0.7325 \pm 0.0086}$ | $0.5645 \pm 0.0082$ | $0.5711 \pm 0.0068$ |
| | 1000 | $\mathbf{0.7570 \pm 0.0073}$ | $0.5605 \pm 0.0063$ | $0.5738 \pm 0.0068$ |
| 50% | 100 | $0.5038 \pm 0.0395$ | $0.5697 \pm 0.0204$ | $\mathbf{0.5990 \pm 0.0199}$ |
| | 200 | $0.5954 \pm 0.0202$ | $0.5683 \pm 0.0117$ | $\mathbf{0.5962 \pm 0.0159}$ |
| | 300 | $\mathbf{0.6197 \pm 0.0224}$ | $0.5702 \pm 0.0102$ | $0.5848 \pm 0.0108$ |
| | 400 | $\mathbf{0.6479 \pm 0.0101}$ | $0.5646 \pm 0.0088$ | $0.5863 \pm 0.0086$ |
| | 500 | $\mathbf{0.6550 \pm 0.0129}$ | $0.5731 \pm 0.0078$ | $0.5878 \pm 0.0070$ |
| | 1000 | $\mathbf{0.6844 \pm 0.0055}$ | $0.5669 \pm 0.0057$ | $0.5815 \pm 0.0054$ |
| 90% | 100 | $0.5225 \pm 0.0739$ | $0.5163 \pm 0.0259$ | $\mathbf{0.5958 \pm 0.0148}$ |
| | 200 | $0.5582 \pm 0.0430$ | $0.5717 \pm 0.0175$ | $\mathbf{0.5978 \pm 0.0191}$ |
| | 300 | $0.5829 \pm 0.0276$ | $0.5736 \pm 0.0136$ | $\mathbf{0.6008 \pm 0.0114}$ |
| | 400 | $\mathbf{0.5981 \pm 0.0268}$ | $0.5715 \pm 0.0128$ | $0.5864 \pm 0.0120$ |
| | 500 | $\mathbf{0.6162 \pm 0.0189}$ | $0.5717 \pm 0.0084$ | $0.5855 \pm 0.0070$ |
| | 1000 | $\mathbf{0.6452 \pm 0.0155}$ | $0.5683 \pm 0.0064$ | $0.5878 \pm 0.0051$ |
